# Dorsal Motor Vagal Neurons Can Elicit Bradycardia and Reduce Anxiety-Like Behavior

**DOI:** 10.1101/2023.11.14.566855

**Authors:** Misty M. Strain, Nicholas J. Conley, Lily S. Kauffman, Liliana Espinoza, Stephanie Fedorchak, Patricia Castro Martinez, Maisie E. Crook, Maira Jalil, Georgia E. Hodes, Stephen B. G. Abbott, Ali D. Güler, John N. Campbell, Carie R. Boychuk

## Abstract

Cardiovagal neurons (CVNs) innervate cardiac ganglia through the vagus nerve to control cardiac function. Although the cardioinhibitory role of CVNs in nucleus ambiguus (CVN^NA^) is well established, the nature and functionality of CVNs in dorsal motor nucleus of the vagus (CVN^DMV^) is less clear. We therefore aimed to characterize CVN^DMV^ anatomically, physiologically, and functionally. Optogenetically activating cholinergic DMV neurons resulted in robust bradycardia through peripheral muscarinic (parasympathetic) and nicotinic (ganglionic) acetylcholine receptors, but not beta-1-adrenergic (sympathetic) receptors. Retrograde tracing from the cardiac fat pad labeled CVN^NA^ and CVN^DMV^ through the vagus nerve. Using whole cell patch clamp, CVN^DMV^ demonstrated greater hyperexcitability and spontaneous action potential firing *ex vivo* despite similar resting membrane potentials, compared to CVN^NA^. Chemogenetically activating DMV also caused significant bradycardia with a correlated reduction in anxiety-like behavior. Thus, DMV contains uniquely hyperexcitable CVNs capable of cardioinhibition and robust anxiolysis.

## Introduction

Cardiovagal neurons (CVNs) send axonal projections from hindbrain to cardiac ganglia through the vagus nerve to elicit cardioinhibitory (i.e., slowing of heart rate; HR) action at rest and during critical cardiorespiratory homeostatic reflexes. Although in mammals most CVNs are found in the nucleus ambiguus (CVN^NA^), 20% of CVNs arise from a second region, the dorsal motor nucleus of the vagus (CVN^DMV^) ^1-6^. While the ability of CVN^NA^ to control chronotropy (e.g. HR) is well characterized ^7-9^, whether this ability extends to CVN^DMV^ remains controversial, despite their extensive innervation of cardiac tissue ^10^. CVN^DMV^ innervate cardiac tissue with unmyelinated, C fibers ^11^, the selective activation of which causes a bradycardia (or a decrease in HR) in multiple mammalian species ^12,13^. In addition, studies of CVN ontogenesis implicate DMV, not NA, as the primary vagal nucleus in lower-order vertebrates and in early mammalian embryonic development, since CVN^NA^ migrate out of DMV ^14,15^, making it possible that CVN^DMV^ retain functional cardioinhibitory activity in mammals.

Despite these anatomical studies, functional studies remain conflicting. Some argue that direct electrical stimulation of DMV does not change chronotropy ^7,16^, while others demonstrate local activation (chemical, electrical, optogenetic, or chemogenetic) elicits bradycardias even if only modestly ^17-21^. However, no activation technique in awake animals used to date rules out incidental stimulation of either neighboring nucleus tractus solitarius (NTS) neurons or inhibitory interneurons within DMV. Stimulation of this latter population could significantly dampen the impact of cholinergic premotor neuron stimulation. Thus, whether cholinergic CVN^DMV^ are capable of controlling chronotropy remains an important but open question.

A recent resurgence of interest in understanding of vagal physiology is driven by the growing number of therapeutic applications for vagus nerve stimulation ^22^. Of particular interest, vagus nerve stimulation is a promising treatment for anxiety and post-traumatic stress disorder ^23,24^. In rats, stimulating the vagus nerve accelerates extinction of conditioned fear ^25,26^ and decreases anxiety-like behavior in the elevated plus maze paradigm ^23^. In humans, vagal nerve stimulation may even reduce treatment-resistant anxiety disorders ^27^. However, the mechanism behind the anxiolytic effect of vagal stimulation is currently unknown. Since the vagus nerve is a bidirectional nerve, carrying sensory information from viscera to brain and motor information from brain to viscera, the extent to which the anxiolytic effect of vagus nerve stimulation depends on sensory or motor signaling is unclear. One possibility is that activating the vagus nerve decreases anxiety by slowing HR, in line with the James-Lange theory of emotions ^28^. Accordingly, a recent study in mice demonstrated that tachycardia (e.g. increase in HR) is sufficient to induce anxiety-like behavior ^29^.

Controversy over the role of CVN^DMV^ may stem in part from differences in CVN^NA^ and CVN^DMV^ electrophysiology and thus their roles in regulating HR. Notably, CVN^NA^ are largely quiescent *in vivo* and *ex vivo* ^30,31^, responding solely to integrated synaptic signaling. While limited data exist on the electrophysiological properties of CVN^DMV^ specifically, other DMV neuronal populations have unique pace-making properties and maintain a relatively high activity *ex vivo* ^32^, which allows for more complex signal processing. One investigation into their activity also suggests that, unlike CVN^NA^, CVN^DMV^ do not demonstrate robust respiratory burst patterning ^33^. Therefore, it is tempting to speculate that CVN^DMV^ integrate and communicate central information differently from CVN^NA^ and represent a functionally unique circuit with respect to CVN^NA^, which is consistent with evidence that CVN^DMV^ and CVN^NA^ target different cardiac ganglia ^34^. A more extensive characterization of the electrophysiological properties of CVN^DMV^ will help build a foundation for understanding their cardioregulatory role, and importantly, for harnessing the therapeutic potential of vagus nerve stimulation. This is critical given that synaptic input to CVN^DMV^ is uniquely sensitive to perturbation of the cardiovascular system, relative to CVN^NA^ ^35^.

The aim of the present study was to characterize the ability of DMV neurons to regulate HR functionally, physiologically, and pharmacologically. We hypothesized that activating choline acetyltransferase positive (Chat+) CVN^DMV^ elicits robust bradycardias. Moreover, we sought to characterize the electrophysiological properties of CVN^DMV^ to determine whether they are distinct from the more extensively studied CVN^NA^. Testing of these hypotheses was accomplished using a combination of opto- and chemogenetic stimulation specifically in Chat+ neurons, pharmacology techniques to determine the regulatory role of CVN^DMV^ on chronotropy, and retrograde labeling paired with electrophysiology procedures to examine differences between the two CVN populations. Our results raise the possibility that CVN^DMV^ play a role distinct from CVN^NA^ in regulating HR.

## RESULTS

### Optogenetic stimulation of Chat+ DMV can elicit bradycardia

To determine if CVN^DMV^ activation is capable of cardioinhibition, we used optogenetics to activate cholinergic neurons in DMV. Specifically, we crossed Ai32 mice expressing channelrhodopsin-2/EYFP fusion protein (ChR2) to loxP floxed choline acetyltransferase (Chat^cre^) mice to selectively express ChR2 in Chat+ neurons (Chat^cre^;ChR2). Chat^cre^;ChR2 and Chat^cre^ mice (Figure 1A) were implanted with HR telemetry devices and fiber optic probes above the right DMV. HR was examined before and during photostimulation. A preliminary study found that 20 Hz (−261 ± 32 bpm; n = 4 mice) elicited a significantly larger bradycardia compared to 5 (−58 ± 28 bpm; repeated measure one-way ANOVA with Šídák’s post-hoc, *p* = 0.0003), 10 (−126 ± 26 bpm; *p* = 0.0064), and 40 (−43 ± 16 bpm; *p* = 0.0002) Hz stimulation to DMV in Chat^cre^;ChR2 mice (data not shown). We further verified that DMV neurons were continuously activated by 20 Hz photostimulation using whole-cell patch-clamp in brain slices from Chat^cre^;ChR2 mice (before: 2.5 ± 1.4 Hz versus during: 13.0 ± 2.7 Hz; repeated measure one-way ANOVA with Tukey’s post-hoc, *p* = 0.0482; n = 5 neurons from two mice; Figure 1B). DMV neurons from Chat^cre^;ChR2 mice also had lower firing frequencies immediately following stimulation (after: 1.4 ± 1.2 Hz; *p* = 0.0321), and they qualitatively demonstrated a pause immediately following stimulation termination that lasted for 10.3 ± 3.9 s with a range of 0.7 to 18 s.

**Figure 1:**
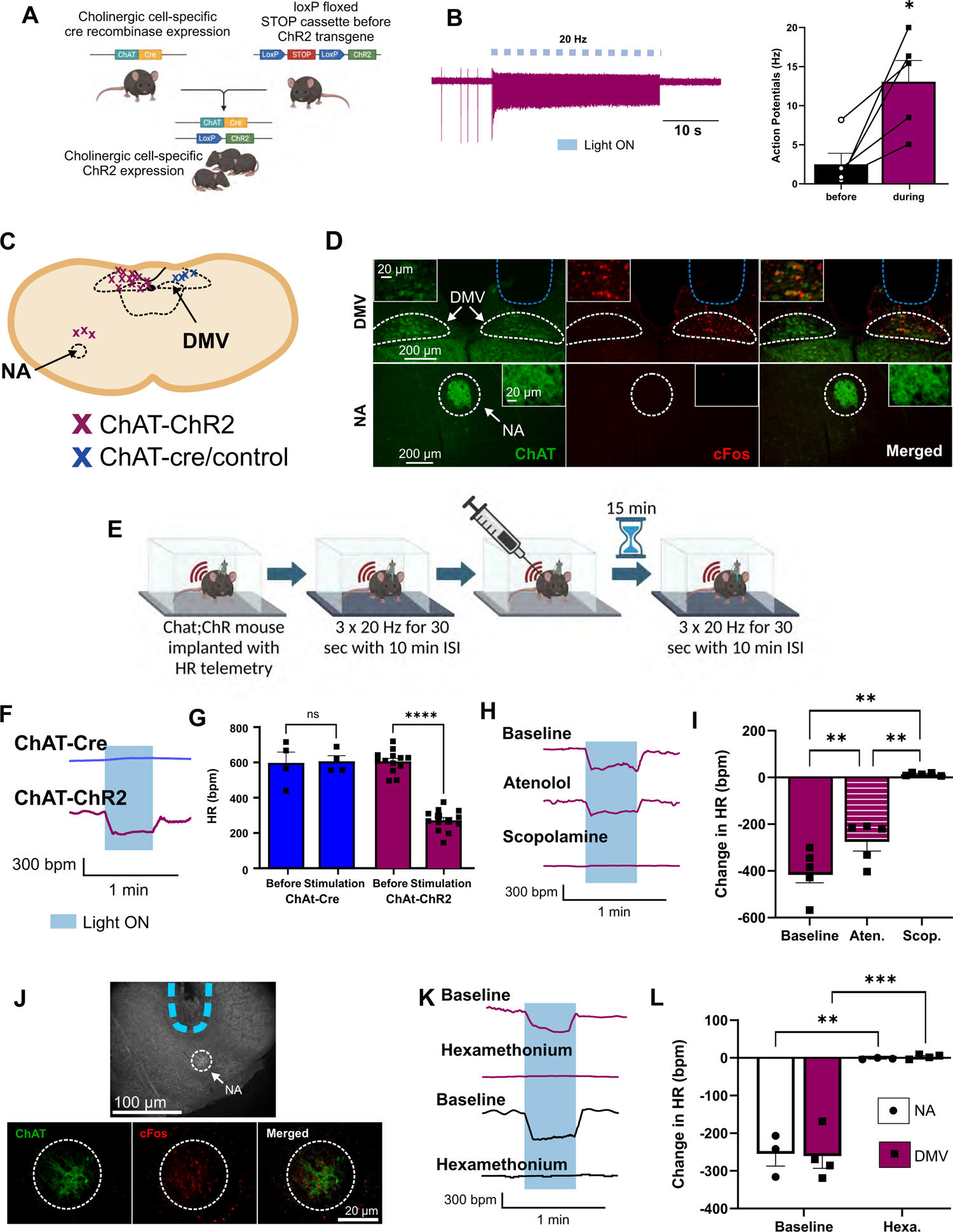
Optogenetic stimulation of DMV elicits bradycardia in male and female mice. Genetic crossbreeding paradigm used to generate transgenic mice harboring loxP-flanked ChR2 in Chat-positive motor neurons (A). Photostimulation-evoked action potentials from DMV motor neurons pace to 20 Hz stimulation (B). Representative diagram illustrating confirmed locations of optogenetic probes (C). Representative images of DMV (top) and NA (bottom) stained for the neuronal activation marker, c-Fos (middle panel; red) and GFP (left panel; green) immunoreactivity confirming c-Fos activation in DMV, but not NA, after DMV photostimulation (D). Schematic illustrating time course of optogenetic studies in awake mice (E). Representative trace (F) and mean HR (G) showing optogenetic stimulation of DMV produced a bradycardia in mice expressing Chat^cre^;ChR2 but not in Chat^cre^ mice. Representative trace (H) and mean HR (I) showing *i.p.* muscarinic parasympathetic blocker, methyl-scopolamine, eliminated the photostimulation-induced bradycardia, while sympathetic blockage with β-1 receptor blocker atenolol mildly reduced this bradycardia. Representative images of probe site (top) and immunohistochemical staining of NA showing c-Fos activation after stimulation of NA (J). Representative trace (K) and mean HR (L) showing the nicotinic antagonist, hexamethonium *(i.p.)* abolished photostimulation-induced bradycardia in both DMV and NA. Bars represent mean and SEM. Blue bars/shading indicates light stimulation. *p ≤ 0.05, **p ≤ 0.01, ***p ≤ 0.001, ****p ≤ 0.0001.

Photostimulating DMV in awake male and female Chat^cre^;ChR2 mice significantly affected HR compared to Chat^cre^ mice. During photostimulation, Chat^cre^;ChR2 decreased HR (271 ± 17 bpm) compared to HR before stimulation (605 ± 18 bpm; repeated measure two-way ANOVA with Šídák’s post-hoc, *p* < 0.0001; n = 13 mice; Figure 1F, G). Photostimulation failed to significantly affect HR during stimulation in control, Chat^cre^ mice (before: 597 ± 61 bpm vs. during: 606 ± 31 bpm, *p* = 0.9633; n = 4 mice), confirming that photostimulation-induced bradycardia in Chat^cre^;ChR2 was not a consequence of off-target effects or exposure to a laser (e.g., tissue necrosis, light diffusion, temperature effects). Finally, photostimulation-induced bradycardia in Chat^cre^;ChR2 was significantly different from HR responses to light in Chat^cre^ control (comparing both mouse lines during stimulation: *p* < 0.0001).

In contrast to photoexcitation of DMV neurons, photoinhibition of cholinergic neurons in DMV of awake mice using Chat^cre^ mice crossed to Ai39 mice conditionally expressing halorhodopsin (NpHR^EYFP^), produced no significant HR response to light (before: 629 ± 45 bpm vs. during: 627 ± 38 bpm; two-tailed paired Student’s t-test, *p* = 0.8950, n = 3 mice). While these results suggest that under these conditions DMV neurons are sufficient to regulate HR but not necessary for control of resting HR, the decay kinetics of NpHR^EYFP^ expressed by Ai39 mice are sufficiently fast enough that it might not cause a robust neural inhibition ^36^.

Since the axons of CVN^NA^ travel dorsally to converge with axons from CVN^DMV^ before turning and exiting the brainstem (e.g., ^37^), it was possible that cardioinhibitory actions of DMV resulted from stimulating nearby CVN^NA^ axons. To rule out CVN^NA^ axonal stimulation, we first examined c-Fos expression in ChAT+ neurons two hours post-DMV stimulation (Figure 1D). Qualitatively robust c-Fos was seen on the stimulated (right) side of DMV with no c-Fos expression in NA. We also confirmed using immunofluorescence that stimulating NA directly causes c-Fos expression (Figure 1J). Finally in a subset of anesthetized Chat^cre^;ChR2 mice, photostimulation of the exposed vagus nerve produced no change in HR (before: 422 ± 61 bpm vs. during: 419 ± 60 bmp; two-tailed paired Student’s t-test, *p* = 0.9723; n = 3 mice). Taken together, axonal expression of ChR2 in NA was not sufficient to activate NA and elicit bradycardia.

### DMV photostimulation-induced bradycardia works through canonical vagal circuits

To confirm that DMV photostimulation affects HR though autonomic pathways, male Chat^cre^;ChR2 mice were administered an intraperitoneal (*i.p.)* injection of hexamethonium to block all nicotinic acetylcholine receptor (nAChR) communication between preganglionic and cardiac-projecting postganglionic parasympathetic neurons prior to DMV (n = 4) or NA (n = 3) photostimulation (Figure 1E). Photostimulation of DMV (−261 ± 32 bpm) produced as similar of a robust bradycardia as photostimulation of NA (−255 ± 32 bpm) (repeated measure two-way ANOVA*, p* = 0.8157). As expected, pretreatment of hexamethonium abolished photostimulation-induced bradycardias regardless of regions (HR responses in DMV: 3 ± 3 bpm and HR responses in NA: −2 ± 1, *p* < 0.0001; Figure 1J-L), and HR was also not different between DMV and NA during photostimulation after pre-treatment with hexamethonium (p = 0.9931).

To confirm that DMV photostimulation affects HR though muscarinic acetylcholine receptor (mAChR) activation from postganglionic parasympathetic neurons to cardiac tissue, consistent with canonical cardiovagal signaling, Chat^cre^;ChR2 mice received *i.p*. injections of scopolamine methylbromide (to block parasympathetic activity) and atenolol (to block sympathetic activity) in a randomized order prior to photostimulation (Figure 1E). In this subset of mice (n = 5), optogenetic activation of DMV neurons again induced robust bradycardia (−405 ± 46 bpm) (Figure 1H, I). Although administration of atenolol modestly reduced DMV light-induced bradycardias (−276 ± 39 bpm; repeated measure one-way ANOVA with Tukey’s post-hoc, *p* = 0.003; Figure 1H, I), atenolol also significantly decreased resting HR (before: 665 ± 21 vs. atenolol: 527 ± 10, *p* = 0.0093; n = 5). Using a simple linear regression, there was a significant negative relationship between resting HR and DMV light-induced bradycardia (*R^2^* = 0.4741, *p* = 0.0277). Therefore, it is likely that the impact of atenolol on DMV-related bradycardia is through overall reductions in resting HR, and not stimulation of postganglionic sympathetic pathways. Unlike atenolol, however, administering scopolamine methylbromide abolished DMV light-induced bradycardias (14 ± 3 bpm, *p* = 0.002, Figure 1H. I). Thus, both nAChR and mAChR activity is required for photostimulation-induced decreases in HR from DMV similar to NA. Some reports suggest that DMV-mediated vagal activity on cardiac tissue occurs through non-canonical nAChR communication between postganglionic parasympathetic neurons to cardiac tissue ^38^. However, since mAChR antagonism abolished DMV light-induced bradycardias, this confirms that DMV neurons induce robust bradycardia through the canonical cardiovagal pathway nAChR→mAChR signaling (and not nAChR→nAChR).

### DMV innervates cardiac tissue through the vagus nerve

Retrograde tracing with rhodamine and cholera toxin subunit B (CT-B) was done in male C57BL6/J mice to calculate the number of CVN^DMV^ and CVN^NA^ labeled by each tracer (n=6-8 mice per tracer for each CVN group; Figure 2A, B). Although both tracers identified positive neurons in NA and DMV, there were fewer traced neurons in CVN^DMV^ (31 ± 19 neurons) compared to CVN^NA^ regardless of tracer (99 ± 16 neurons; repeated measure two-way ANOVA with Šídák’s post-hoc, *p* < 0.0001; Figure 2C). In all animals, both right and left DMV contained positive labeling. In one animal examined for right versus left distribution, the right DMV contained more retrograde labeling (68 neurons) compared to the left DMV (45 neurons). This pattern was similar to previous reports in rats ^39^. In addition, CT-B (47 ± 36 neurons) labeled significantly fewer neurons than rhodamine regardless of region (83 ± 33 neurons; *p* = 0.0472). There was not a significant interaction between CVN location and tracer (*p* = 0.8294). To confirm that retrograde labeling of CVN^DMV^ required an intact vagus nerve, a unilateral left vagotomy prior to cardiac injection of rhodamine was performed in a separate group of animals (n = 3 mice, Figure 2D). Unilateral vagotomy eliminated labeling of CVN^DMV^ ipsilateral to vagotomy (2 ± 1 labeled CVN^DMV^) compared to the contralateral side (32 ± 8 labeled CVN^DMV^; two-tailed paired Student’s t-test, *p* = 0.0483; Figure 2E). Taken together, DMV contains neurons that retrogradely label from cardiac tissue in a vagus-dependent manner.

**Figure 2:**
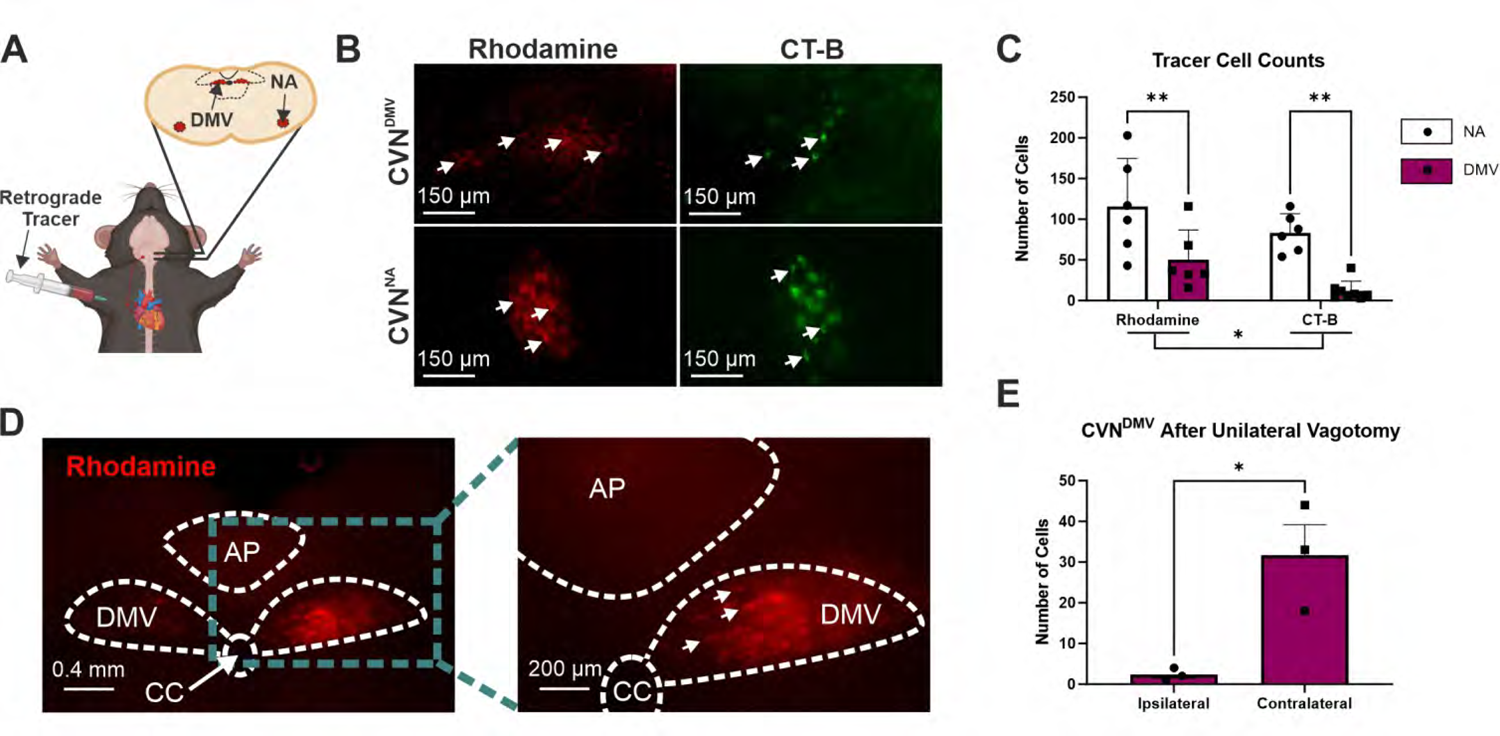
DMV innervates cardiac tissue through the vagus nerve in male mice. Schematic showing injection into the epicardial fat pad and labeling in both CVN brain regions (A). Representative images of DMV and NA after cardiac injection of retrograde tracers, rhodamine (left in red) and cholera toxin subunit B (CT-B; right in green) (B). Both rhodamine and CT-B significantly labeled cardiac projecting neurons in NA and DMV (C). Representative images of DMV after a right cervical vagotomy (C). Representative image (D) and mean cell count (E) showing a right cervical vagotomy significantly attenuated CVN^DMV^ numbers ipsilateral to vagotomy. Bars represent mean and SEM. *p ≤ 0.05, **p ≤ 0.01, ***p ≤ 0.001, ****p ≤ 0.0001.

### CVN^DMV^ show spontaneous firing and have larger input resistance *ex vivo* compared to CVN^NA^

To characterize the electrophysiological properties of CVN^DMV^ in relation to CVN^NA^, additional studies examined the general excitability of CVNs in male C57BL6/J mice using whole-cell patch-clamp electrophysiology under current clamp configuration. Retrogradely labeled CVNs were identified using visual inspection for rhodamine and anatomical landmarks (Figure 3A). Post-hoc biocytin recovery confirmed location and cholinergic phenotype (Figure 3B). CVN^DMV^ exhibited more spontaneous firing (8/14 neurons from 10 mice) compared to CVN^NA^ (0/10 neurons from 9 mice; Mann Whitney test, *p* = 0.0064), which were completely devoid of any spontaneous firing activity (Figure 3C-D). Of CVN^DMV^ exhibiting spontaneous firing (57%), the average frequency was 1.28 ± 0.31 Hz.

**Figure 3:**
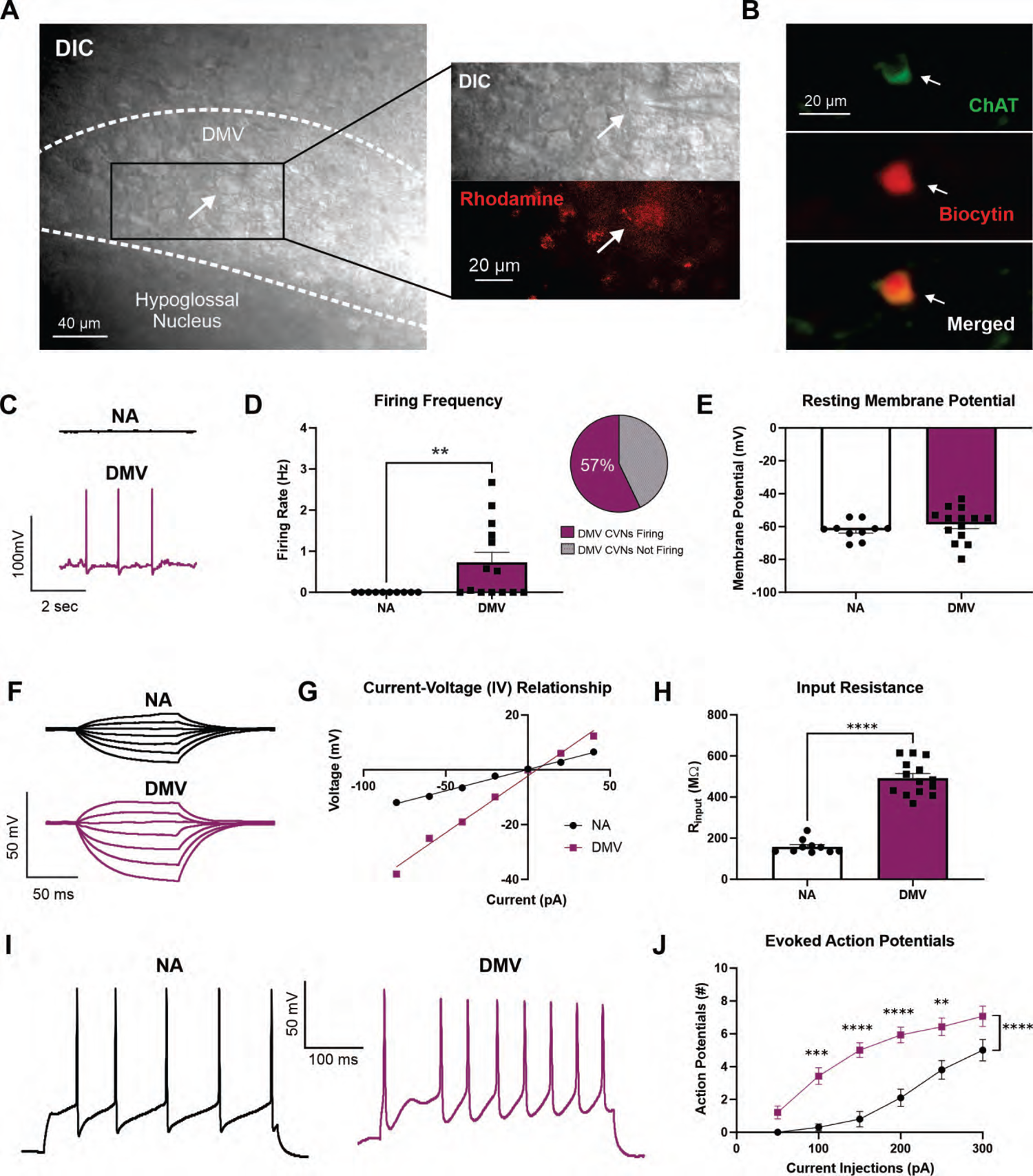
Differential electrophysiological properties of CVN^DMV^ compared to CVN^NA^ in male mice. Rhodamine-positive DMV neurons showing pipette (top) and rhodamine (bottom; red) (A). Representative immunofluorescence image of CVN^DMV^ showing biocytin recovered patched cardiac-labeled neurons are cholinergic (B). Representative trace (C) and mean firing rate (D) of CVN^DMV^ neurons show significantly higher spontaneous firing rates compared to CVN^NA^, with the majority of CVN^DMV^ firing. No statistical differences in resting membrane potential were found between CVN^DMV^ and CVN^NA^ (E). Representative traces of membrane responses from CVN^NA^ (top) and CVN^DMV^(bottom) to stepped current injections (F). Current-voltage (I-V) relationship graph obtained from CVN^NA^ and CVN^DMV^ (G). R_input_ was higher in CVN^DMV^ compared to CVN^NA^ (H). Representative action potential responses in CVN^NA^ (top) and CVN^DMV^ (bottom) in response to 300 pA injection of direct depolarizing current (I). Action potential response curves were higher in CVN^DMV^ compared to CVN^NA^ in response to 50 pA-step injections of direct depolarizing current (J). *p ≤ 0.05, **p ≤ 0.01, ***p ≤ 0.001, ****p ≤ 0.0001.

No statistical differences in resting membrane potential were found between CVN^DMV^ (−58.60 ± 2.72 mV; n = 14 neurons from 10 mice) and CVN^NA^ (−62.18 ± 1.75 mV; n = 10 neurons from 9 mice; two-tailed unpaired Student’s t-test, *p* = 0.3256, Figure 3E). However, we found that CVN^DMV^ (492.00 ± 22.02 MΩ; n = 14 neurons from 10 mice) demonstrated significantly larger input resistance (R_input_) than CVN^NA^ (158.20 ± 10.61 MΩ; n = 10 neurons from 9 mice; two-tailed unpaired Student’s t-test, *p* < 0.0001; Figure 3F-H). Additionally, CVN^DMV^ (30.0 ± 1.5 pF) were significantly smaller in size than CVN^NA^ (61.1 ± 6.6 pF; two-tailed unpaired Student’s t-test, *p*<0.0001) based on their recorded capacitance.

### CVN^DMV^ are hyperexcitable compared to CVN^NA^ *ex vivo*

To determine if there were any differences in excitability between CVN^DMV^ and CVN^NA^, CVNs were examined for general excitability through stepped depolarizations of current (Figure 3I). As expected, regardless of neuronal type, stepped current injections evoked a current dependent increase in action potential frequency (repeated measure two-way ANOVA with Šídák’s post-hoc, *p* <0.0001; Figure 3J). Additionally, a significant effect of neuronal type regardless of current injection was found, with CVN^DMV^ exhibiting a significantly greater number of evoked action potentials (5 ± 1 number of action potentials, n = 14 neurons from 10 mice) compared to CVN^NA^ (2 ± 1 number of action potentials, n = 10 neurons from 9 mice; *p* < 0.0001). Finally, we found a significant interaction between current injection and neuronal type (*p* < 0.0001), with significantly more evoked action potentials in CVN^DMV^ compared to the CVN^NA^ at 100 (0.30 vs. 3.42 ± 0.5; *p* = 0.0001), 150 (0.80 vs. 5.00 ± 0.5; *p* < 0.0001), 200 (2.10 vs. 5.93 ± 0.5; *p* < 0.0001), and 250 pA (3.80 vs. 6.43 ± 0.6; *p* = 0.0017); Figure 3J). Taken together, CVN^DMV^ are more excitable than CVN^NA^.

### Chemogenetic activation of DMV suppresses HR for up to 8 hours

Although our optogenetic studies could rule out interneuron activity, they could not exclude the possibility that off target stimulation of nearby Chat+ neurons (e.g., hypoglossal neurons) affected HR. Therefore, to further confirm that activating DMV neurons specifically decreases HR, we used an alternative strategy, chemogenetically activating DMV neurons with the excitatory designer receptor, hM3Dq. Here, hM3Dq was specifically targeted to DMV neurons through an intersectional genetics approach which leverages the co-expression of *Chat* and *Phox2b* genes by DMV neurons but not by neighboring neurons (i.e., hypoglossal) ^40,41^. First, to validate this intersectional approach, Chat^Cre^;Phox2b^Flp^ mice were crossed with R26^ds-HTB^ mice, a reporter line which expresses nuclear-localized, green fluorescent protein (GFP)-tagged histone 2b protein (H2b-GFP) upon recombination by both Cre and Flp. In the resulting Chat^Cre^;Phox2b^Flp^;R26^ds-HTB^ mice, essentially all peripherally-projecting DMV neurons, as labeled by a systemic (*i.p*.) retrograde tracer, Fluorogold, were H2b-GFP immunofluorescent (136/139 DMV neurons from four mice (97.8%) were both Fluorogold+ and H2b-GFP+; representative image in Supplemental Figure 1A). These results confirm that Chat^Cre^ and Phox2b^Flp^ are co-expressed by all DMV vagal efferent neurons and thus validate our intersectional approach.

Next, to express hM3Dq specifically in DMV neurons, an adeno-associated virus (AAV) in which Cre- and Flp-dependently expresses a hemagglutinin (HA)-tagged hM3Dq (AAV1-CreON/FlpON-hM3Dq-HA; Figure 4A) was injected into DMV of male and female Chat^Cre^;Phox2b^Flp^;R26^ds-HTB^ mice. This resulted in hM3Dq-HA expression in the vast majority of H2b-GFP+ DMV neurons (“DMV-hM3Dq” mice; Figures 4A-C). Importantly, we failed to detect any hM3Dq-HA expression in NA (Figure 4B-C) or DMV of mice lacking Cre and Flp recombinases (Supplemental Figure 1B), confirming the specificity of our injection strategy and AAV expression.

**Figure 4:**
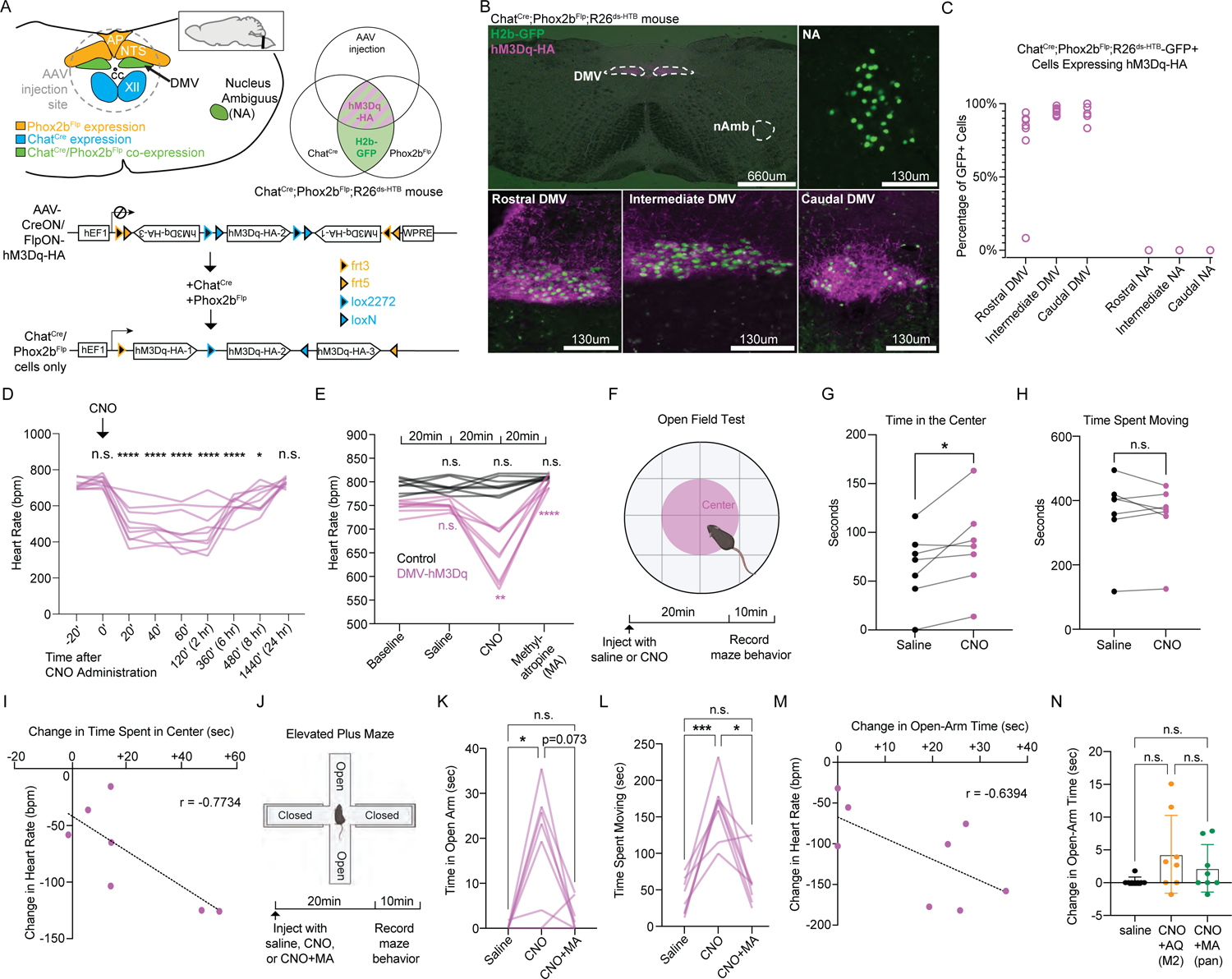
Chemogenetic stimulation of DMV produces bradycardia and reduces anxiety in both male and female mice. Illustration of coronal hindbrain section showing injection site and *Chat* and *Phox2b* expression (A, top left); Schematic of AAV1-CreON/FlpON-hM3Dq-HA viral construct (A, bottom); Venn diagram showing expected expression of hM3Dq-HA and H2b-GFP (A, top right). Representative images of rostral, intermediate, caudal DMV and NA stained for HA (magenta) and H2b-GFP (green) immunoreactivity in Chat^Cre^;Phox2b^Flp^;R26^ds-HTB^ mice after DMV injection of AAV1-CreON/FlpON-hM3Dq-HA (B). Percentage of H2b-GFP+ cells immunoreactive for hM3Dq-HA in the rostral, intermediate and caudal DMV and NA (C). Effect of CNO on HR in DMV-hM3Dq mice over 24-hour period. CNO (1mg/kg, *i.p*.) injected at time = 0 min (D). Effect of *i.p.* saline, CNO and CNO+MA on HR in DMV-hM3Dq (magenta) and control mice (black); comparisons are to baseline (E). Schematic of open field experiment (F). Effects of *i.p.* saline vehicle and CNO on time in the center of the open field, in seconds (G). Correlation between changes in HR and center time between saline and CNO conditions in DMV-hM3Dq mice (H).Schematic of elevated plus maze (EPM) experiment (I). Effects of *i.p.* saline vehicle, CNO and CNO+MA administration on time spent moving in the elevated plus maze, in seconds (J). Effect of *i.p.* saline vehicle, CNO and CNO+MA administration on open-arm time in the elevated plus maze, in seconds (K). Effect of *i.p.* saline vehicle, CNO and CNO+AQ administration on open-arm time in the elevated plus maze, in seconds. (L). Correlation between changes in HR and open-arm time between saline and CNO conditions in DMV-hM3Dq mice (M). Effects of *i.p.* MA and AQ-RA-741 on open-arm time in elevated plus maze, compared to saline; same saline, CNO, and CNO+MA data as in Figure 4K (N). *p ≤ 0.05, **p ≤ 0.01, ***p ≤ 0.001, ****p ≤ 0.0001.

To determine whether chemogenetically activating DMV neurons affects HR, hM3Dq ligand clozapine N-oxide (CNO; 1 mg/kg; *i.p*.) was administered while measuring HR by non-invasive electrocardiography (ECG; n=8 mice per genotype). Administering CNO to DMV-hM3Dq mice significantly decreased HR for up to 8 hours compared to baseline (baseline: 718 ± 10 bpm; 20 min: 497 ± 34 bpm, *p* = 0.0012; 40 min: 497 ± 30 bpm, *p* = 0.0008; 60 min: 467 ± 36 bpm, *p* = 0.0007; 2hr: 469 ± 39 bpm, *p* = 0.0011; 6hr: 598 ± 19 bpm, *p* = 0.0006; 8hr: 637 ± 24 bpm, *p* = 0.0274; Figure 4D). HR returned to baseline by 24 hours post-CNO (24hr: 731 ± 9 bpm, *p* = 0.7247). As expected, the acute bradycardia after CNO administration required peripheral muscarinic signaling, as it was abolished by administering the peripheral muscarinic blocker, methyl-atropine bromide (MA; 1.0 mg/kg; *i.p*.), 20min after CNO (baseline: 747 ± 5 bpm; after CNO: 640 ± 19 bpm *vs*. baseline, *p* = 0.0078; after MA: 801 ± 4 bpm *vs*. baseline, *p* = 0.0004; Figure 4E). On the other hand, administering CNO to hM3Dq-negative control mice failed to significantly alter HR (p > 0.05 *vs*. baseline or vehicle; Figure 4E; Supplemental Figure 1C), indicating that CNO alone did not affect HR. These results, together with our optogenetics studies, provide robust evidence that DMV neurons can decrease HR through peripheral muscarinic signaling.

### Activating DMV neurons reduces anxiety-like behavior

Tachycardia increases anxiety-like behavior in mice ^29^, consistent with the James-Lange theory that physiological cues can drive emotional states ^28^. We therefore wondered whether the CNO-induced bradycardia in DMV-hM3Dq mice corresponded to any change in anxiety-like behavior. To investigate, male and female DMV-hM3Dq mice and control mice (n=8 per group) were administered CNO or saline vehicle and then assessed behaviorally by the open field test, an assay which measures anxiety-like behavior based on time spent in the center of an open arena (Figure 4F) ^42,43^. Strikingly compared to vehicle, CNO treatment in DMV-hM3Dq mice significantly increased their time spent in the center of the open field (CNO: 85.31 ± 17.38 s vs. Vehicle: 64.55 ± 13.98 s; two tailed paired Student’s t-test, *p* = 0.0379, Figure 4G). The increased time spent in the center was not due to an increase in overall motility since total time spent moving did not differ significantly between saline and CNO treatments (two tailed paired Student’s t-test, *p* = 0.4643, Figure 4H). In addition, CNO’s effects on HR and center time in DMV-hM3Dq mice were moderately correlated (R^2^ = 0.7734, *p* = 0.0414; Figure 4I). Thus, activating DMV neurons caused a correlated decrease in HR and anxiety-like behavior in the open field test.

To confirm the change in anxiety-like behavior, we assessed another cohort of DMV-hM3Dq mice in a different measure of anxiety, the elevated plus maze ^44^. CNO treatment significantly increased the time DMV-hM3Dq mice spent in the open-arms of an elevated plus maze, relative to vehicle treatment, indicating a decrease in anxiety-like behavior (CNO: 16.9 ± 4.83 s vs. Vehicle: 0.23 ± 0.23 s, repeated measure ANOVA with Tukey’s post-hoc, *p* = 0.0235; Figure 4K). CNO treatment also significantly increased other signs of anxiolysis in DMV-hM3Dq mice: time spent moving (CNO: 162.6 ± 14.4 s vs. Saline: 40.7 ± 8.7 s; *p* = 0.0006; Figure 4L); head dip events (CNO: 4.149 ± 1.022 dips vs. Saline: 0.2414 ± 0.1424 dips; *p* = 0.0259; Supplemental Figure 1E); and average velocity (CNO: 2.339 ± 0.3068 cm/s vs. Saline, 0.6606 ± 0.2317 s, *p* = 0.0011; Supplemental Figure 1F). Importantly, however, treating hM3Dq-negative control mice with CNO did not affect their open-arm time, indicating that CNO itself does not decrease anxiety-like behavior (*p* = 0.3868; Supplemental Figure 1D). In addition, co-administering MA (“CNO+MA”) to DMV-hM3Dq mice largely prevented CNO’s effects on measures of anxiolysis, indicating that anxiolysis, as with HR, require peripheral mAChR signaling (Figure 4K; Supplemental Figure 1E, F). Specifically, compared to vehicle treatment, CNO+MA failed to significantly affect open-arm time, time spent moving, velocity or head dip events, relative to treatment with a saline vehicle (p < 0.05; Figure 4K, L, Supplemental Figure 1D,E). As with our open field test, the decrease in HR was trended towards a moderate correlation with the increase in open-arm time (R^2^ = 0.6394, *p* = 0.0878; Figure 4M). Thus, our results show that activating DMV neurons similarly decreases HR and anxiety-like behavior, and that these effects each depend on peripheral muscarinic signaling.

To identify the muscarinic receptors mediating anxiolysis, we repeated our HR and elevated plus maze studies on DMV-hM3Dq mice but used a selective inhibitor of muscarinic type 2 receptors (M2), AQ-RA 741 (“AQ”) ^45^, which does not appear to cross the blood-brain barrier ^46^. Administering AQ after CNO reversed the CNO-induced bradycardia (repeated measure ANOVA with Tukey’s post-hoc, *p* = 0.0016; Supplemental Figure 1G). Importantly, in contrast to CNO alone, administering CNO with AQ (CNO+AQ) did not significantly affect time spent in the open-arms relative to saline treatment (*p* = 0.2560; Figure 4N). These results suggest that activating DMV neurons decreases anxiety-like behavior through a M2-dependent mechanism.

## DISCUSSION

The present study characterized CVN^DMV^ anatomically, physiologically, and functionally. Retrograde tracing from the cardiac fat pad confirmed the presence of CVN in DMV using two different tracers, rhodamine and CT-B, and this labeling required an intact vagus nerve. Electrophysiological comparison of retrogradely-labeled CVN^DMV^ and CVN^NA^ indicate distinct circuits, with CVN^DMV^ being significantly more excitable with higher input resistances and spontaneous activity *ex vivo* whereas CVN^NA^ were not. Optogenetically activating DMV neurons caused bradycardia, which was completely abolished by muscarinic and nicotinic antagonism. Finally, intersectional chemogenetic activation of DMV neurons also caused bradycardia and a correlated decrease in anxiety-like behavior, both of which also required peripheral muscarinic signaling, as they were blocked by muscarinic antagonism.

Previous efforts to characterize *in vivo* electrophysiological properties of DMV neurons are limited, possibly a result of their prominent C-fiber phenotypes and long latency conduction times ^33^ making typical antidromic spike identification challenging. Our retrograde labeling findings are in accordance with previous reports, which suggest that only ∼20% of CVN exist within DMV in totality (both sides), making the likelihood of recording from these numbers *in vivo* relatively low ^14^. Importantly, CVNs signal organization is an established example of divergent amplification, meaning that changes in HR arise from modulation (activation or inhibition) of a relatively small number of CVNs^30^. Therefore, whole-cell patch-clamp recordings paired with retrograde labeling in adult animals—albeit confounded by the absence of relevant synaptic inputs *in vivo*—is a powerful technique that future studies should continue to use to aid in the investigation of this population of neurons both in health and disease.

Our *ex vivo* results are consistent with electrophysiological recording in other DMV populations. DMV neurons demonstrate an on-going activity phenotype ^32,47,48^. This was true in the present study as ∼57% of CVN^DMV^ demonstrated a spontaneously active firing pattern during the recording period. With regards to CVNs, this on-going firing activity is unique to CVN^DMV^ since no CVN^NA^ recorded exhibited any spontaneous activity in our slice preparation, which is also consistent with previous results from CVN^NA^ ^31^. While future studies will need to demonstrate how this spontaneous activity is generated, it is consistent with the intrinsic pace-making properties of other DMV neurons. However, it also suggests that CVN^DMV^ may represent a functionally discrete circuit for cardiac-related vagal motor output. Previous studies suggest that CVN^DMV^ have little respiratory-related burst activity, despite robust lung-related afferent activity ^33^, which is in contrast to the CVN^NA^ which show robust respiratory-related burst activity both *in vivo* and *ex vivo* ^9^. While we are only beginning to elucidate the complexity of vagal circuits as it relates to any vagal motor output, there is increasing evidence for distinct circuits related to vagal sensory afferents. Similar distinct circuits also likely exist in vagal motor nuclei ^41,49,50^. While some parallel circuits have converging anatomy and function, they may be distinct in terms of their physiological roles ^50^. Finally, CVN^DMV^ demonstrated a significantly higher input resistance, higher number of spiking activity to current injections, and smaller size compared to CVN^NA^. During optogenetic stimulation, CVN^DMV^s were also silenced for several seconds after stimulation implicating additional electrophysiological properties (i.e., unique ion channel contributions or stimulation-induced plasticity in neurotransmitter contributions) that could be investigated. Therefore, CVN^DMV^ are more excitable than CVN^NA^. In other brain regions, this type of behavior confers a coincidence detection phenotype over a simple integrator ^51^. Therefore, future work should continue to investigate how CVNs in NA and DMV differ in their microcircuit construction in relation to the basic circuit building blocks of critical homeostatic regulating networks.

Despite historical controversy about the capacity of CVN^DMV^ to impact HR, our results demonstrate that optogenetic and chemogenetic activation of Chat+ DMV neurons each caused robust bradycardia (∼56% and ∼65.5% of resting HR, respectively) in awake, behaving mice. Cardioinhibitory responses using optogenetic techniques in urethane-anesthetized rats and ketamine/xylazine-anesthetized mice demonstrated variable strength of the response ^18,21^, implicating differences in anesthesia for differences in effect size. Alternatively, previous species used (namely cats and rats) may exhibit a lesser degree of DMV-activation driven bradycardia than mice because of differences in vagal tonus. Since mice have a lower vagal tone than rats for example ^52-54^, CVN^DMV^ activation in mice may recruit a larger number of CVN^DMV^ which were not previously active. Although it is possible that fast decay kinetics of NpHR^EYFP^ in the Ai39 mouse line limited the ability to significantly inhibit DMV neurons ^36^, the lack of an effect on HR of optogenetic inhibition of DMV in mice is similar to previous reports using inhibitory chemogenetic receptors ^17^. Therefore, while the present study introduced a significant refinement of techniques using Chat^cre^ mice to eliminate the impact of known interneurons within DMV, it may suggest that CVN^DMV^ activity is not necessary for generation of resting HR, similar to previous report ^17^.

While our Chat-targeted approach avoids influence of interneurons within DMV, the anatomical proximity of vagal axons from NA and the optical fiber in DMV ^14^ raises the possibility that DMV stimulation resulted in axonal stimulation of CVN^NA^ neurons. However, we failed to detect c-Fos expression in NA and DMV contralateral to the optical fiber, despite robust c-Fos immunoreactivity in the ipsilateral DMV. In addition, direct photostimulation of the vagus nerve in Chat^cre^;ChR2 animals did not produce bradycardia. Finally, the more specifically targeted chemogenetic receptor hM3Dq expressed in DMV, but not other nearby cholinergic regions such as NA or hypoglossal neurons, caused a similar cardioinhibition to DMV stimulation in Chat^cre^;ChR2. Taken together, these data provide robust evidence that solely activating DMV is sufficient for cardioinhibition (Figure 1D).

As expected, pharmacological testing confirmed that DMV stimulation-induced bradycardia requires mACh receptor activation in mice, since scopolamine (but not atenolol) and MA abolished optogenetically- and chemogenetically-induced bradycardia, respectively. Notably, while the traditional concept of vagal motor signaling to cardiac tissue requires nicotinic receptor-dependent communication between preganglionic and postganglionic parasympathetic neurons, some studies report that intracardiac ganglia harvested from SA node also show mACh receptor-dependent neurotransmitter and calcium mobilization ^55^. Additional studies have attributed pharmacological differences in vagal fiber activation to C-fiber (presumably from DMV) versus B-fiber bradycardia (presumably from NA) since hexamethonium did not block the HR responses to non-myelinated fiber activation in these studies ^56^. Still, it is important to note that the aforementioned studies did not comment on whether postganglionic neurons were isolated from cardiac nodal cells, nor did they directly address the effect of neurotransmitter release on the observed HR responses. Our findings are in accordance with the canonical signaling pathway between preganglionic and postganglionic parasympathetic neurons, as pre-treatment with the mACh receptor antagonist, scopolamine, completely abolished light-induced bradycardia in ChAT^cre^;ChR2 mice.

Recent research suggests that HR can influence emotional states, supporting a theory of emotion separately proposed by the physiologist Carl Lange and psychologist William James over a century ago. As explained by James: “*My thesis […] is that the bodily changes follow directly the Perception of the exciting fact, and that our feeling of the same changes as they occur IS the emotion.”* ^57^. In other words, the James-Lange theory holds that HR does not increase because of anxiety; rather, the perception of HR increase is the anxiety. Consistent with this theory, a recent study in mice demonstrated that optogenetically increasing HR is sufficient to drive anxiety-like behaviors in the elevated plus maze ^29^. Inversely, beta-blockers such as propranolol, which robustly decrease HR ^58^, also reduce anxiety-like behaviors in mice ^59,60^. Beta-blockers and other drugs which decrease HR have shown promise for treating anxiety and related disorders ^61^, though the mechanism is not yet known. Studies of brain activity in humans and mice suggest the insular cortex may play a key role in sensing HR and other interoceptive cues ^29,62-64^ and so could couple the perception of HR to emotional state.

Importantly, however, while our study is the first to our knowledge to demonstrate a correlation between vagally-mediated bradycardia and anxiolysis, more research is needed to establish a causal relationship. Our results with the cardioselective M2 receptor antagonist AQ-RA 741 ^45^, which does not appear to cross the blood-brain barrier ^46^, suggests a cardiogenic mechanism for the anxiolysis we observed when activating DMV neurons. For instance, one possibility is that DMV decreases anxiety-like behavior by decreasing HR. This would agree with recent studies showing that optogenetically increasing HR is anxiogenic ^29^, whereas pharmacologically decreasing HR is anxiolytic^65^. Indeed, the M2 receptors blocked by AQ-RA 741 are enriched in the heart relative to other organs innervated by the DMV ^45,66^ and are necessary for vagally-mediated bradycardia ^67^. However, M2 receptors are also present in the gastrointestinal tract where they interact with M3 receptors to modulate smooth muscle contraction ^68^. Therefore, our results do not rule out the possibility that DMV decreases anxiety-like behavior through its projections to the gut ^68,69^. Further research, e.g., targeting organ-specific DMV circuits, is needed to establish the mechanism by which DMV neurons can decrease anxiety. Whether other physiological functions of DMV, such as gut motility, glucose metabolism, and suppressing inflammation <u>^58^</u>, are involved remains unknown.

In summary, this study demonstrates the existence of CVN^DMV^ neurons and their ability to suppress HR and anxiety-like behavior. The cardioinhibitory and behavioral effects both require mACh receptor activity, and our results also suggest that nACh receptors are involved in the signaling between CVN^DMV^ and cardiac parasympathetic neurons, implicating canonical vagal pathways in DMV’s facilitation of cardioinhibitory activity. CVN^DMV^ also has distinct electrophysiological properties, namely spontaneous firing activity and higher R_input_, compared to CVN^NA^ *in slice*. Therefore, we further speculate that CVN^DMV^ represents a unique population of CVNs, distinct from the more thoroughly characterized CVN^NA^. Further research is needed to identify the molecular profile of CVN^DMV^, the mechanisms underlying their distinct electrophysiological properties, and their physiological role.

## ACKNOWLEDGEMENTS

Funding was provided by a Pathway to Stop Diabetes Initiator Award 1-18-INI-14, an NIH R01 HL153916 to J.N.C., an NIH R01HL157366 to C.R.B. and an NIH T32 HL007446 for M.M.S.

## AUTHOR CONTRIBUTIONS

M.M.S., N.J.C., L.S.K., M.J., S.B.G.A., J.N.C., and C.R.B. designed experiments. M.M.S, L.E., and S.F. performed optogenetic and cardiac tracing studies, including HR analysis, and all histology/immunofluorescence. L.E. and C.R.B. performed electrophysiological studies. A question from G.E.H. inspired the study of anxiety-like behavior. N.J.C., L.S.K., M.E.C., and P.C.M. performed chemogenetic study of HR and anxiety-like behavior. N.J.C. and L.S.K. performed histology and immunofluorescence experiments related to chemogenetic studies. M.M.S., N.J.C., J.N.C., L.E., and C.R.B. prepared figures. M.M.S., N.J.C., L.S.K., C.R.B., and J.N.C. wrote the manuscript with input from all authors.

## DECLARATION OF INTEREST

Authors declare no competing interest.

---

**Supplemental Figure 1:**
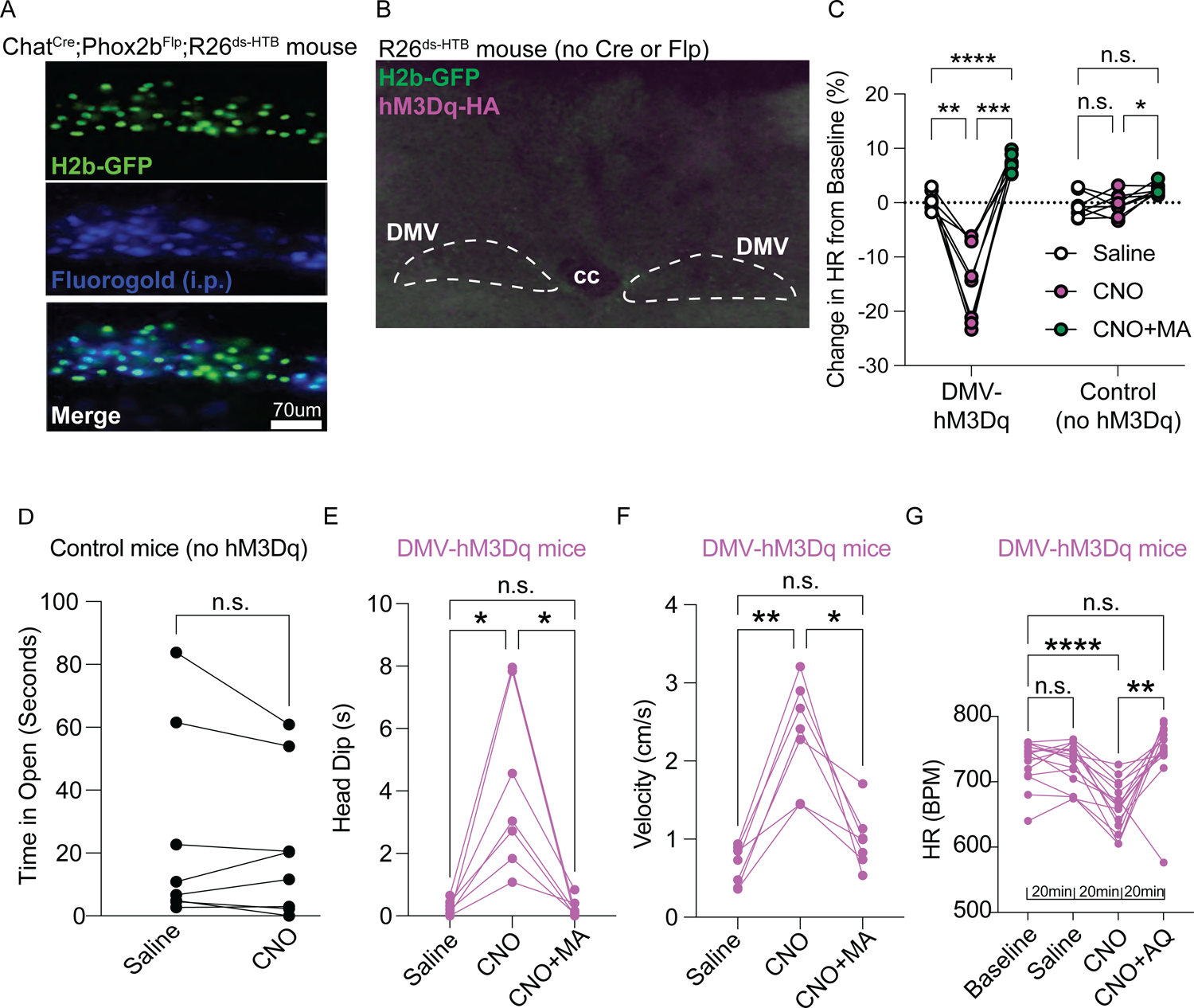
Representative images of intermediate DMV hemisphere stained for H2b-GFP (green) and fluorogold immunoreactivity (blue) in Chat^Cre^;Phox2b^Flp^;R26^ds-HTB^ mice after *(i.p.)* systemic injection of retrograde tracer Flurogold (136 of 139, 97.8%, Fluorogold+ DMV neurons were also H2b-GFP+); n=4 mice) (A). Representative images of intermediate DMV stained for HA (magenta) and H2b-GFP (green) immunoreactivity in R26^ds-HTB^ mice (Cre-/Flp-) after DMV injection of AAV1-CreON/FlpON-hM3Dq-HA (B). Effect of *i.p.* saline (white), CNO (magenta) and CNO+MA (green) on HR in DMV-hM3Dq and control mice; relative to vehicle (C). Effect of saline or CNO on control mice, which lack hM3Dq expression, open-arm time in the elevated plus maze, n=8 mice (D). Effect of saline, CNO and CNO+MA in DMV-hM3Dq mice on head dip events and velocity in the elevated plus maze (E-F). Effect of saline, CNO and CNO+AQ on HR in DMV-hM3Dq (magenta) mice; same experimental design and baseline, saline, and CNO data as in Figure 4E (G). *p ≤ 0.05, **p ≤ 0.01, ***p ≤ 0.001, ****p ≤ 0.0001.

## LEAD CONTACT

Further information and requests for resources and reagents should be directed to and will be fulfilled by the lead contact, Carie R. Boychuk.

## MATERIALS AVAILABILITY

All data reported in this paper will be shared by the lead contact upon request. This study did not generate new unique reagents or report original code.

## Notes

### Competing Interest Statement

The authors have declared no competing interest.

### Summary of Updates

Figures and text revised for clarity; additional behavioral and pharmacological studies added which strengthen the original conclusions.

